# Analysing complex metagenomic data with MicroWineBar

**DOI:** 10.1101/742684

**Authors:** Franziska Klincke, Martin Abel-Kistrup, Sofie Maria Gilberte Saerens, Simon Rasmussen

## Abstract

An important step in metagenomics studies is to identify which species are present in a sample as well as to compare samples from different environments. Here we introduce MicroWineBar, a graphical tool for analyzing and comparing metagenomics samples. MicroWineBar can visualize the abundances of metagenomics samples in line and bar graphs, as well as analyse the richness and diversity. For a PCA as well as a differential abundance analysis, the abundance data is treated as compositional data and center log-ratio transformed. We use MicroWineBar to analyse two different years of wine fermentation as well as data from a human microbiome study of colorectal cancer. Importantly, MicroWineBar does not require any programming skills, is intuitive and user friendly. MicroWineBar is available at https://github.com/klincke/MicroWineBar and as a python package from the Python Package Index.

## Introduction

Metagenomics is the application of sequencing techniques to study the communities of microbial organisms in an environment [1]. Some examples of environments which have been studied include sea water from the Sargasso Sea [2], acid mine drainage [3] and the human gut [4]. One basic question in metagenomics studies is which species are present in a sample from a specific environment and whether there are differences in species composition between environments. However, analysis of metagenomic data is not straightforward due to large numbers of reads that must be processed and that the microbial composition of the samples are often largely unknown. To investigate these questions it is important to explore the relative abundances of species or higher taxonomic ranks. Determining statistically differential abundant species between two environments can be achieved using either nonparametric methods such as the Wilcoxon rank sum test or taking into account that count data from metagenomic studies is compositional. This can be done by the center log-ratio transformation (CLR) which is applied before using for instance ANCOM [5] for differential abundance testing. Additionally, visualization and dimensionality reduction is needed to get an overview of metagenomic samples.

Many bioinformatic tools exist for visualising metagenomic data and Sudarikov *et al.* provide a comprehensive overview [6]. One of the first tools which was developed was MEGAN [7–9]. MEGAN displays the taxonomic hierarchy by node-link diagrams where each node has a small, log-scaled quantitative chart. The advantage of this approach is that each node is represented in the hierarchy. Another tool for visualising relative abundances is Krona [10] where subdivided pie charts display an embedded hierarchy. This is an easy way to display several taxonomic ranks at the same time. However, it is difficult to compare relative abundances of two species since the abundances are represented by angle. Cleveland *et al.* demonstrated that it is easier to compare values which are represented by length (as in a bar graph) than by angle (as in a pie chart) [11]. Further, Keanu is another visualization tool to explore biodiversity in metagenomes that is able to display the hierarchical taxonomy but is only suited for analysing one sample at a time [12]. The above mentioned tools are mainly for visualising metagenomics data with the exception of MEGAN which also offers to compare several samples. Nonetheless, it does not offer log-ratio transformations which are recommended for compositional data such as metagenomics data.

We have developed a new tool for statistical analysis, dimensionality reduction and visualization of metagenomics datasets. The tool is called MicroWineBar and is able to display relative abundances in bar graphs interactively. MicroWineBar not only includes features to compare metagenomic samples but also enables the user to compare two groups of compositional metagenomic samples, e.g. from two environments. It supports common data exploration techniques including Principal Component Analysis (PCA) scatter plots and Shannon diversity index plots to interpret variability in a metagenomic dataset. In addition, it provides both scatter and box plots for analysing the species richness. These plot can be saved as high-resolution figures. We have implemented two methods to determine differentially abundant species between two groups of samples: the nonparametric Wilcoxon rank sum test and a differential abundance test using ANCOM (Analysis of composition of microbiomes) [5]. The latter calculates the pairwise log-ratios between all species and performs a significance test on it to reduce the False Discovery Rate (FDR), from which many differential abundance analysis methods (e.g. the commonly used DESeq2 [13]) suffer. ANCOM treats the count data from metagenomic samples as compositional data because the relative abundances within a sample have to sum to one [14]. Additionally, MicroWineBar can provide information about specific species through both Wikipedia and PubMed in the tool. We exemplify the use of MicroWineBar using metagenomics samples from wine fermentations of Bobal grapes and by re-analysing a published human microbiome dataset [15]. In both examples we point out the differences between two sample groups. In the wine data we compare samples from two years and show that there are differences between the vintages, especially regarding richness and diversity. For the human microbiome dataset we can partly replicate the findings of Zeller *et al.* between the two sample groups. MicroWineBar is available at https://github.com/klincke/MicroWineBar.

## Results

### Overview of MicroWineBar

MicroWineBar is designed to read simple text input files containing estimates for abundances in metagenomics samples. In order to be more flexible, the input of MicroWineBar is not tied to any specific abundance measurement tool. Instead we provide custom scripts to generate abundance tables for the following programs: MGmapper [16], Bracken [17] which is run on top of Kraken [18] and MetaPhlAn [19].

These abundance tables contain relative (and absolute) abundances with taxonomic annotations determined from mapping to whole genome databases, k-mer based databases or marker gene databases, respectively. The design of MicroWineBar follows the visual information-seeking mantra: overview first, zoom, filter and details on demand [20]. These key tasks are recommended for designing advanced graphical user interfaces and will be addressed in the following. The main window of MicroWineBar will display relative abundances in bar graphs. By default, it starts with displaying a bar graph of the relative abundances of the species present in the first sample which was loaded. Each species is represented by one bar and its height corresponds to the relative abundance of the species in that sample (Fig 1). One can change the taxonomic rank, filter out individual species or taxonomic groups or filter for a minimum (relative) abundance. Additionally, information about the species can be retrieved in the form of a Wikipedia summary or PubMed publications. One can create line and stacked bar graphs for several samples on all taxonomic ranks. In addition, one can also compare two groups of samples, e.g. to identify differentially abundant species. We demonstrate this in the following paragraphs using two example datasets.

**Fig 1.**
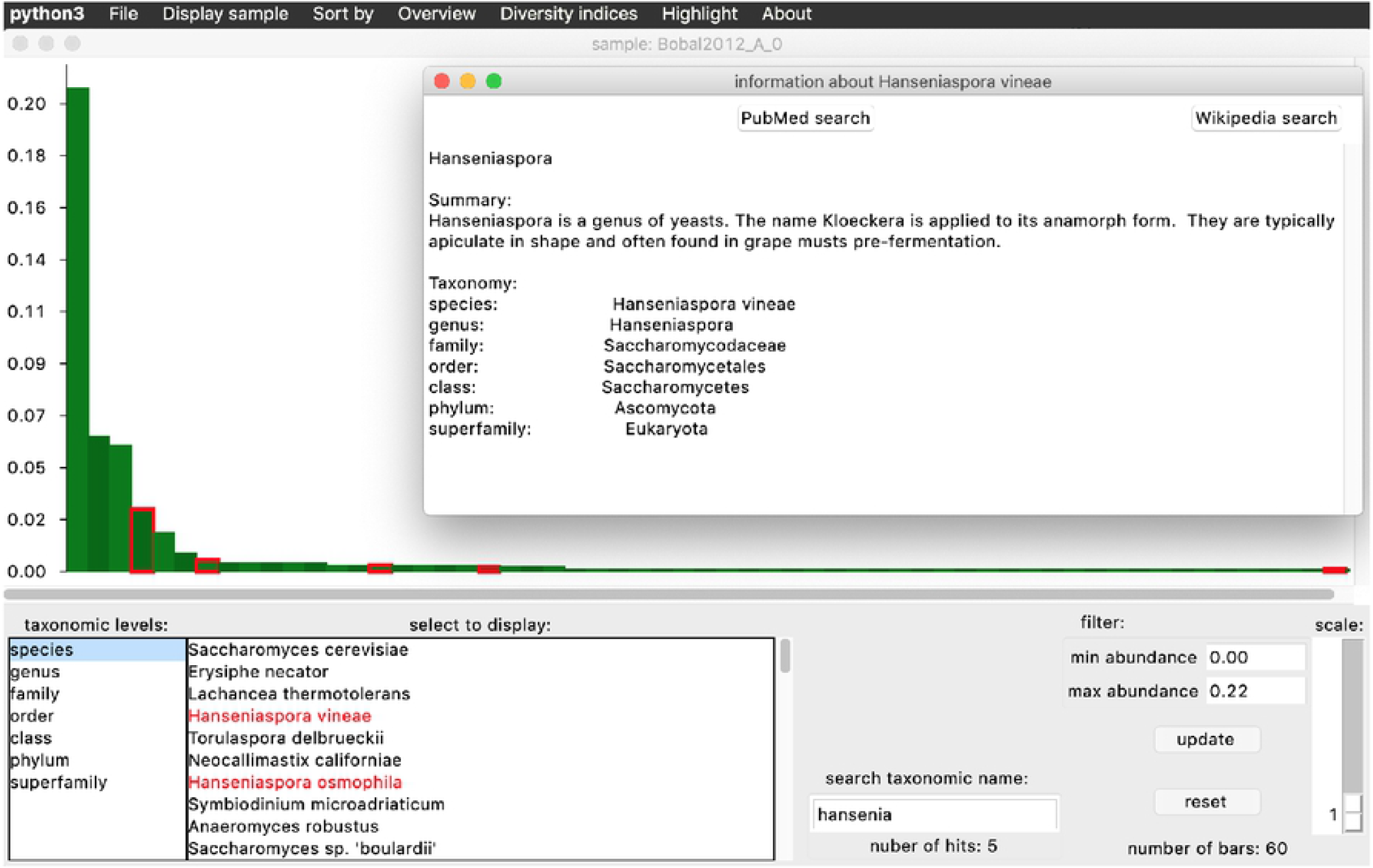
Main window of MicroWineBar. An example of the main window of MicroWineBar where one metagenomic wine sample is displayed. All hansenia* species are highlighted in red and a popup window with information for the species *Hanseniaspora vineae* is displayed. One can see a summary from Wikipedia, the taxonomy as well as links to PubMed and Wikipedia.

### Differences in the wine fermentation of two vintages

We tested MicroWineBar using a dataset of 21 metagenomics samples from two different years of wine fermentations from a winery in the La Mancha region in Spain. The aim was to identify differential abundant species between the fermentations of Bobal grapes from 2012 (6 samples from one fermentation tank with a mean of 17,285,292 reads per sample) and from 2013 (15 samples from two fermentation tanks with a mean of 13,514,142 reads per sample). In the PCA analysis the samples from the two years were well separated into two groups (Fig 2A) with the variance explained by the first two principal components as 28.72% and 16.05%, respectively. When analysing the species richness we found that samples from 2012 had a significantly lower richness compared to the samples from 2013 (p-value: 8e-10) with a median of 87 compared to 334 (Fig 2B). Additionally, we found the Shannon diversity index to be significantly higher (p-value: 6e-2) for the 2013 samples (median of 2.15) compared to the 2012 samples (median of 0.57) (Fig 2C).

**Fig 2.**
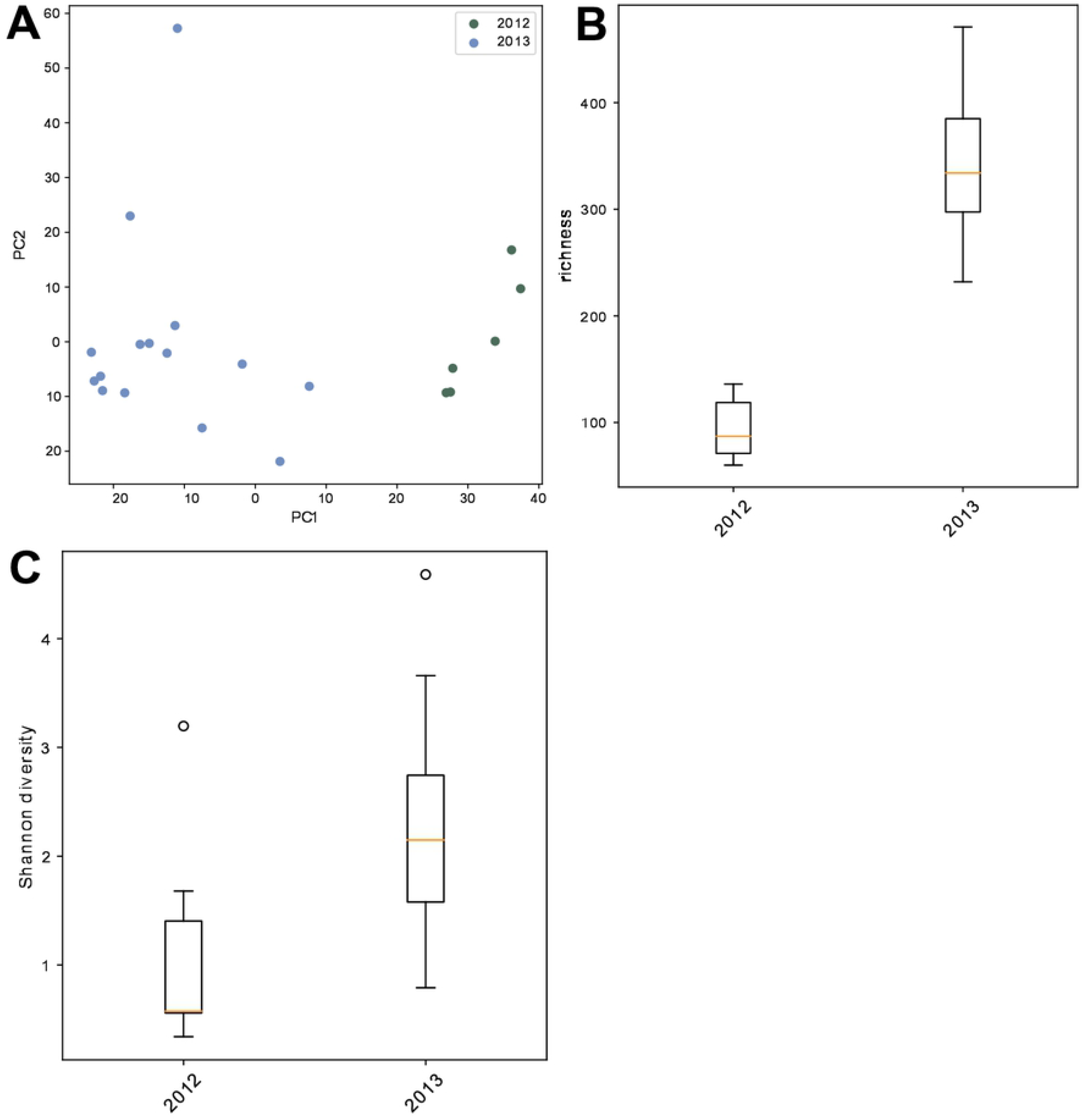
Diversity of metagenomic wine samples from Bobal fermentations. Wine samples from 2012 are compared to wine samples from 2013. A: The Principal Component Analysis (PCA) scatter plot shows a clear separation of the two vintages on the first Principal Component (PC). B: Boxplot of the species richness where the 2012 samples show a significantly lower richness. C: Boxplot of the Shannon diversity showing that the 2013 samples are more diverse. For the boxplots the lower and upper hinges correspond to the first and third quartiles and the median is indicated with an orange line. The upper and lower whiskers extend up to 1.5 * interquartile range (IQR). Outliers are represented by circles.

Using the conservative ANCOM method we obtain ten species to be differentially abundant. Of these Aspergillus niger, Bradyrhizobium sp. BTAi1, Penicillium expansum, Pseudoalteromonas prydzensis, Pseudomonas cerasi, Pseudomonas syringae, *Rhodanobacter glycinis* and *Starmerella bacillaris* are more abundant in 2013 whereas *Hanseniaspora vineae* and *Candida sp. LDI48194* are more abundant in 2012 (Table 1). The presence of *H. vineae* is in good concordance with previous findings that at the beginning of wine fermentation microorganisms originating from the grapes such as *Candida, Pichia, Brettanomyces, Aureobasidium* or *Hanseniaspora* [21] are present. Some of these are considered spoilage microorganisms such as *Brettanomyce*s [20], however later during the alcoholic fermentation the principal wine yeast *Saccharomyces cerevisiae* dominates the community.

**Table 1.** Differential abundance analysis of the wine samples. ANCOM was used to obtain the results. Only the significantly differential abundant species are displayed. ANCOM calculates the pairwise log ratios between all species and performs a one-way ANOVA (Analysis of Variance) test to determine if there is a significant difference in the log ratios with respect to the species of interest. “W” is then the W-statistic which is the number of pairwise log ratios where the species is significantly different between the two years. Also, the median and maximum abundance are reported for both years.

| Species | W | Median abundance |  | Maximum abundance |  |
| --- | --- | --- | --- | --- | --- |
|  |  | 2012 | 2013 | 2012 | 2013 |
| *Aspergillusniger* | 844 | 1.03e-06 | 4.22e-03 | 1.38e-04 | 3.33e-02 |
| *Bradyrhizobiumsp.BTAi1* | 875 | 1.03e-06 | 2.75e-02 | 1.30e-06 | 2.41e-01 |
| *Candidasp.LDI48194* | 875 | 2.13e-04 | 1.30e-06 | 6.09e-03 | 1.30e-06 |
| *Hanseniasporavineae* | 799 | 8.26e-04 | 6.10e-05 | 5.78e-02 | 5.08e-04 |
| *Penicilliumexpansum* | 845 | 1.30e-06 | 9.43e-04 | 1.30e-06 | 1.07e-02 |
| *Pseudoalteromonasprydzensis* | 840 | 1.30e-06 | 1.43e-03 | 1.30e-06 | 6.57e-03 |
| *Pseudomonascerasi* | 852 | 1.30e-06 | 2.17e-03 | 1.30e-06 | 1.18e-02 |
| *Pseudomonassyringae* | 875 | 1.30e-06 | 2.29e-02 | 1.30e-06 | 1.06e-01 |
| *Rhodanobacterglycinis* | 825 | 1.30e-06 | 7.30e-06 | 1.30e-06 | 1.90e-02 |
| *Starmerellabacillaris* | 830 | 1.30e-06 | 6.75e-04 | 1.30e-06 | 2.08e-03 |

### Replicating a human microbiome study

In addition to investigating metagenomic samples from wine fermentations we used MicroWineBar to re-analyse previously published data. Here we investigated 141 human gut microbiome samples of colorectal carcinoma (CRC) patients and tumor-free controls [15]. In the original study the authors created a classifier based on 22 species to distinguish CRC patients from controls based on the microbiome profiles and taxonomic markers. We wanted to replicate their findings of taxonomic markers, namely two *Fusobacterium* species, *Porphyromonas asaccharolytica* and *Peptostreptococcus stomatis* enriched in the CRC samples. To generate count data to be analyzed in MicroWineBar we mapped the reads to reference genome databases and determined species abundance using MGmapper [16]. Hereafter we loaded the count data into MicroWineBar for analysis. Initially we performed a PCA analysis to identify differences between the two groups (Fig 3A), however we were not able to identify any clear differences between the phenotypes. We next investigated species richness and found it to be slightly higher in the CRC samples compared to the control with a median of 199 and 193.5, respectively. However this was not statistically significant (p-value: 6e-2) (Fig 3B). Compared to this the Shannon diversity index was slightly higher (p value: 1e-1) in the control samples (median of 5.43) compared to the CRC samples (median of 5.24) (Fig 3C). Both richness and diversity were also not found to be significantly different in the study by Zeller *et al.* [15]. We found *Fusobacterium* species to be present in some CRC and in none of the control samples. Additionally, we found *P. asaccharolytica* to be present in 75% of the CRC samples compared to only 60% of the control samples. Finally, many *Bacteroides* species were present in all CRC as well as most of the control samples. *Bacteroides* are a major component of the human gut microbiota. To investigate differential abundance between the two groups we applied the conservative ANCOM method. This resulted in only two species being differentially abundant, namely *P. stomatis* and *Parvimonas micra* that were both more abundant in the CRC samples (Table 2).

**Fig 3.**
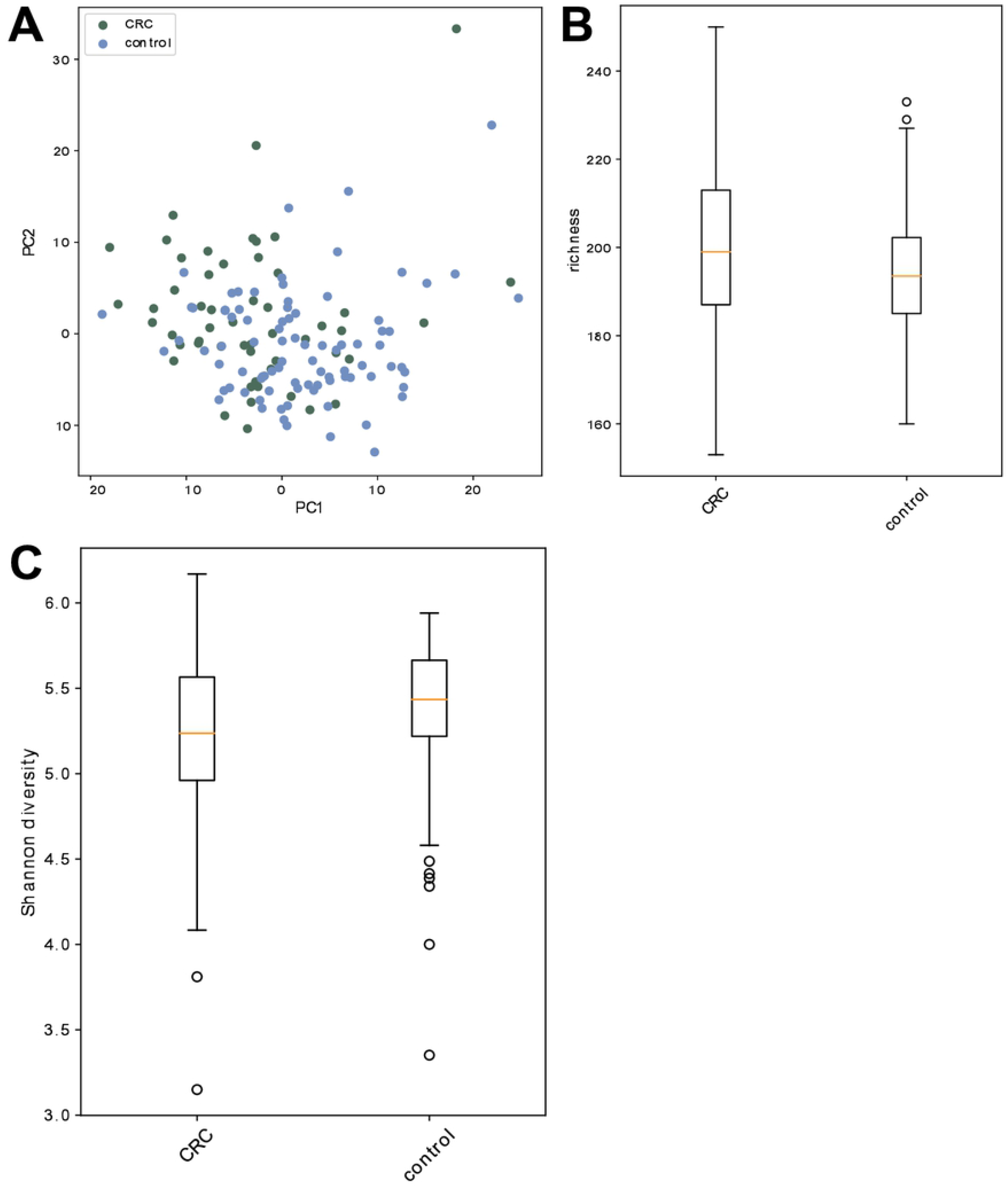
Re-analysing a human microbiome dataset with colorectal cancer (CRC) and control samples. A: A Principal Component Analysis (PCA) scatter plot displaying the CRC (green) and the control (blue) samples. B: Boxplot of the species richness showing no significant difference between the CRC and the control samples. C: Boxplot of Shannon diversity showing no significant difference between the CRC and the control samples. For the boxplots the lower and upper hinges correspond to the first and third quartiles and the median is indicated with an orange line. The upper and lower whiskers extend up to 1.5 * interquartile range (IQR). Outliers are represented by circles.

**Table 2.** Differential abundance analysis of human microbiome samples. ANCOM was used to obtain the results. Only the significantly differential abundant species are displayed. ANCOM calculates the pairwise log ratios between all species and performs a one-way ANOVA (Analysis of Variance) test to determine if there is a significant difference in the log ratios with respect to the species of interest. “W” is then the W-statistic which is the number of pairwise log ratios where the species is significantly different between the two years. Also, the median and maximum abundance are reported for the sample groups.

| Species | W | Median abundance |  | Maximum abundance |  |
| --- | --- | --- | --- | --- | --- |
|  |  | CRC | control | CRC | control |
| *Parvimonasmicra* | 373 | 3.08E-05 | 5.72E-06 | 7.81E-02 | 1.31E-03 |
| *Peptostreptococcusstomatis* | 410 | 5.72E-06 | 5.72E-06 | 1.36E-02 | 6.23E-05 |

## Discussion

MicroWineBar is designed to be a generic visualisation tool for metagenomic samples. This means that it is not tied to a specific analysis toolkit for preparing the input. Among other features it displays relative abundances of species of one or several samples in bar graphs without showing the hierarchy of the taxonomic classifications. Additionally, one can compare groups of samples. For this the data is log-ratio transformed. As expected there are more wine spoilage organisms in the wine fermentation samples from 2013 compared to 2012 and with this an increased species richness in general. This was likely due to differences in weather conditions, where humid conditions favor spoilage organisms such as *Hanseniaspora, Aspergillus, Botrytis* and *Aureobasidium* [21, 22]. The year of 2013 was generally a humid year in Europe and therefore a challenging year for winemaking (Fig 4B). In the La Mancha region, the average humidity was significantly (p-value 0.049) higher in 2013 compared to 2012, especially in the growing season from April to October (p-value 0.001). This is also reflected in that generally wines from the La Mancha region were graded much higher (excellent) in 2012 compared to 2013 (good) [23].

**Fig 4.**
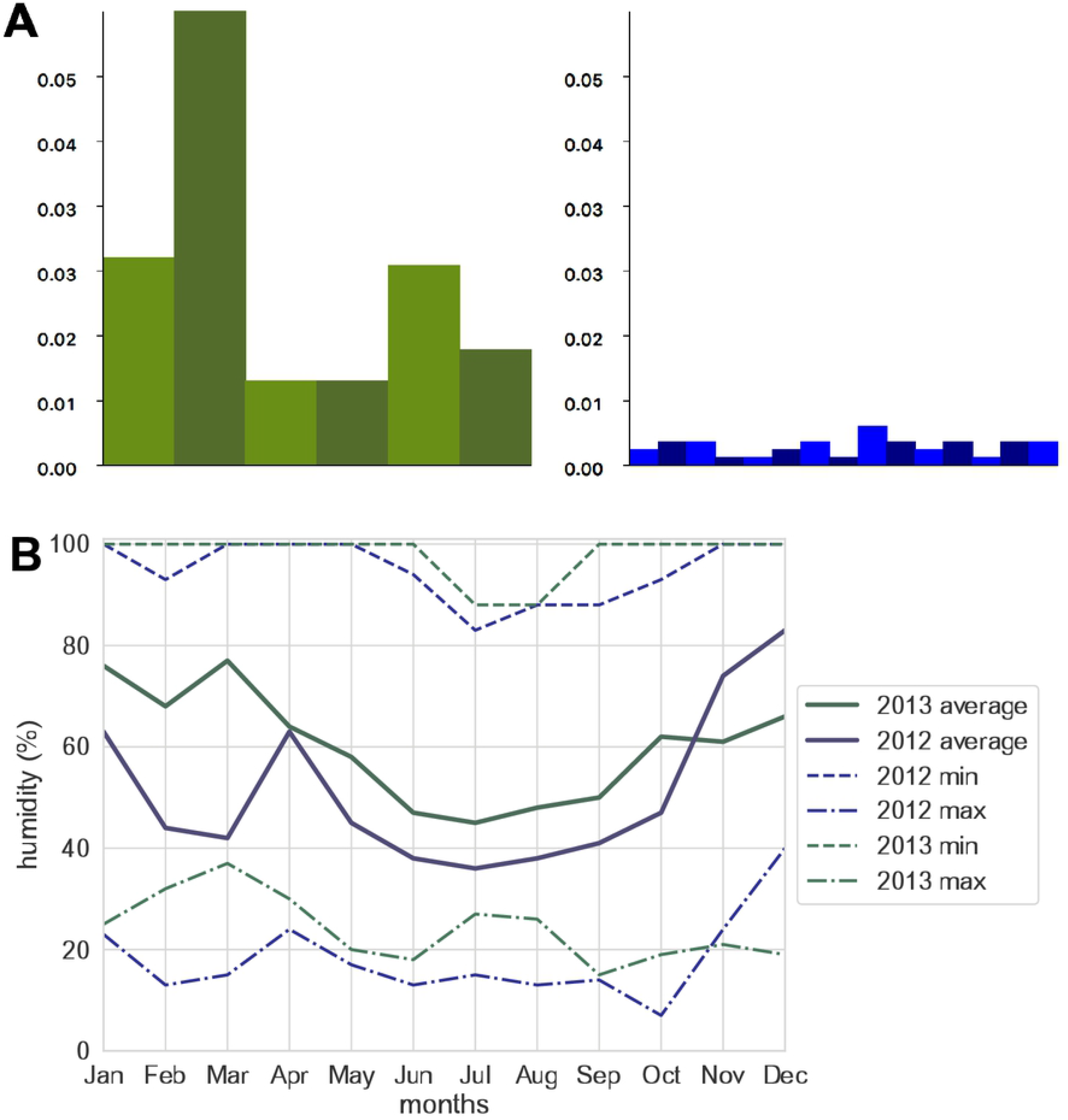
Comparing metagenomic wine samples from Bobal fermentations of 2012 with samples from 2013. A: Abundances of *Hanseniaspora vineae* in 2012 (left) and 2013 (right). The labels on the x-axis indicate the order in which the samples were taken during the fermentation. The 2013 samples were taken from two fermentation tanks. B: Average humidity in La Mancha, Spain, in 2012 (blue) and 2013 (green). The average humidity in La Mancha was significantly higher (p-value: 0.049) in 2013 compared to 2012. The range of the humidity for both years is indicated by broken lines. Data from [24].

Non-*Saccharomyces* wine yeasts were in the past considered as spoilage yeasts but have recently attracted more attention as they are believed to positively modify the wine aroma [22, 25, 26]. The fact that the non*-Saccharomyces* yeast *H. vineae* is more abundant in 2012 than in 2013 as well as that in 2012 it is more abundant than *H. uvarum* is interesting. This might have an influence on the wine aroma for the two years even though the relative abundance of *H. vineae* was low. *Candida sp. LDI48194* belongs to the family *Debaryomycetaceae* and is only lowly abundant in 2012 and absent in 2013. Furthermore *S. cerevisiae* is clearly dominating the fermentations from both years and was even more abundant in the 2013 samples compared to the 2012 samples. For the human microbiome dataset, we identified with ANCOM only one of the two species which contribute most to the classifier in the study by Zeller *et al.* to be differentially abundant as well, namely *P. stomatis.* Additionally, we identified *P. micra* as being differentially abundant which is in concordance with other studies [27]. This means that we could only partially replicate the results. This might be due to the fact that we used another program to align the reads to different databases to get the taxonomic annotations. In other words we might have started with a different set of species found in the samples which of course would influence the result of an analysis. In addition, this might be due to the fact that we used ANCOM which takes into account that microbiome data is compositional. This also points out a problem with metagenomics in general, namely the reproducibility of results from metagenomic studies [28]. MicroWineBar enables researchers without any programming skills to perform analysis and visualization of complex metagenomics datasets. We hope that MicroWineBar will contribute to making analysis of compositional metagenomics data more accessible for non-bioinformaticians.

## Materials and methods

### Implementation of MicroWineBar

MicroWineBar is written in Python3 (v.3.6.8) and uses the GUI package Tkinter. It needs to be installed locally as a Python package and runs on both Mac OS and Linux. The pandas (v.0.23.4) [29] package is used to represent the data in data frames. The scikit-bio (v.0.5.3) package is used to calculate the Shannon diversity index, the Bray-Curtis dissimilarity, to create the PCoA and to perform the differential abundance test using ANCOM [5]. The scipy (v.1.1.0) [30] package was used for the nonparametric Wilcoxon rank sum test. In general, the Benjamini-Hochberg p-value correction was used for multiple hypothesis testing and a corrected p-value of less than 0.05 was considered statistically significant. The matplotlib (v.2.2.3) [31] package was used to create figures. Welch’s unequal variances t-test from the scipy package was used to test for statistical difference between two sample groups, i.e. for richness and Shannon diversity. A p-value of less than 0.05 was considered statistically significant. The module wikipedia (v.1.4.0) was used to retrieve the summary of a Wikipedia entry and the module webbrowser was used to open a website in a browser. MicroWineBar takes files with absolute (and relative) abundances and taxonomic annotations created by Bracken [17], MetaPhlAn [19] and MGmapper [16] as input. For the latter output files from several databases need to be merged with the mgmapper2microwinebar.py script. Scripts are provided to prepare the input.

### Processing of wine samples

The 21 shotgun metagenomic Bobal wine samples were obtained from two studies. From Melkonian *et al.* we only used the Bobal samples from the two control tanks [32]. The sampling points for the six samples from 2012 were 0h, 16h, 24h, 32h, 48h and 96h after fermentation. After 24h the tank was inoculated with S. cerevisiae to start the alcoholic fermentation. The sequencing reads from both years were generated in the following way. For DNA isolation, cells were pelleted from 50 ml of wine centrifuged at 4,500 g for 10 minutes and subsequently washed three times with 10 ml of 4°C phosphate buffered saline. The pellet was mixed with G2-DNA enhancer (Ampliqon, Odense, Denmark) in 2 ml tubes and incubated at room temperature for 5 min. Then 1 ml of lysis buffer (20 mM Tris-HCl- pH 8.0, 2 mM EDTA and 40 mg/ml lysozyme) was added to the tube and incubated at 37°C for one hour. An additional 1 ml of CTAB/PVP lysis buffer (50) was added to the lysate and incubated at 65°C for one hour. DNA was purified from 1 ml of lysate with an equal volume of phenol-chloroform-isoamyl alcohol mixture 49.5:49.5:1 and the upper aqueous layer was further purified with a MinElute PCR Purification kit and the QIAvac 24 plus (Qiagen, Hilden, Germany), according to manufacturer’s instructions, before it was eluted in 100 ul DNase-free water. Prior to library building, genomic DNA was fragmented to an average length of 400 bp using the Bioruptor XL (Diagenode, Inc.), with the profile of 20 cycles of 15 s of sonication and 90 s of rest. Sheared DNA was converted to Illumina compatible libraries using NEBNext library kit E6070L (New England Biolabs) and blunt-ended library adapters. The libraries were amplified in 25-ml reactions, with each reaction containing 5 μl of template DNA, 2,5 U AccuPrime Pfx Supermix (Invitrogen, Carlsbad, CA), 1X Accuprime Pfx Supermix, 0.2 uM IS4 forward primer and 0.2 uM reverse primer with sample specific 6 bp index. The PCR conditions were 2 minutes at 95°C to denature DNA and activate the polymerase, 11 cycles of 95°C for 15 seconds, 60°C annealing for 30 seconds, and 68°C extension for 40 seconds, and a final extension of 68°C for 7 minutes. Sequence data for the 2012 has been deposited at ENA with the accession number ERP111084 and the sequence data for 2013 is available at ENA with the accession number ERP112039. The raw reads were trimmed and quality-filtered with cutadapt (v. 1.16) [33]. The first 10 bp were removed from the reads, the phred quality cutoff was set to 20 and the minimum length to 30. To get the taxonomic annotation the remaining reads were then mapped with MGmapper (v. 2.7) to the following databases: VitisVinifera, Plant, Human, Bacteria, Bacteria_draft, MetaHitAssembly, HumanMicrobiome, Fungi, FungiDB, Brettanomyces, NewWine, Archaea, Virus, Protozoa, Plasmid, Invertebrates. The databases VitisVinifera, FungiDB, Brettanomyces and NewWine were custom databases containing the grape genome and genomes of species which are especially interesting with respect to wine fermentation. MGmapper was run with the option to remove PCR duplicates turned on, the maximum edit distance was set to 0.05 and the minimum number of Matches+Mismatches for a valid read was set to 30. Then the python script mgmapper2microwinebar.py was run for each sample to merge the output of MGmapper so that each sample consists of only one file and can be imported in MicroWineBar. In MicroWineBar the following phyla were filtered: *Annelida, Arthropoda, Chordata, Mollusca*, *Nematoda*, *Platyhelminthes* and *Streptophyta*. The reason for this is that we are only interested in the microorganisms but still wanted to know from which other organisms DNA was found in the samples.

### Processing of colorectal cancer dataset

The shotgun metagenomic dataset analyzed here was downloaded from ENA (ERP005534) [15] and all 88 control and 53 CRC shotgun metagenomic paired end samples were processed. The reads were trimmed and quality-filtered with the same settings as the wine samples. To get the taxonomic annotation the reads were then also mapped with MGmapper [16] with the same settings as the wine samples (v. 2.7) to the following databases: Human (Human reference sequence GRCh38.p12) and HumanMicrobiome [4]. Hereafter the python script mgmapper2microwinebar.py was run for each sample to merge the output of MGmapper. In MicroWineBar the reads that mapped to human were filtered out since we are interested in the differentially abundant microorganisms between the CRC and the control samples.

## Acknowledgements

This study was funded by the Horizon 2020 Programme of the European Commission within the Marie Skłodowska-Curie Innovative Training Network “MicroWine” (grant number 643063). SR was supported by the Novo Nordisk Foundation grant NNF14CC0001.

